# Information-based rhythmic transcranial magnetic stimulation accelerates learning during auditory working memory training

**DOI:** 10.1101/2024.01.05.572158

**Authors:** Heather T. Whittaker, Lina Khayyat, Jessica Fortier-Lavallée, Megan Laverdière, Carole Bélanger, Robert J. Zatorre, Philippe Albouy

## Abstract

Rhythmic transcranial magnetic stimulation (rhTMS) has been shown to enhance auditory working memory manipulation, specifically by boosting theta oscillatory power in the dorsal auditory pathway during task performance. It remains unclear whether these enhancements i) persist beyond the period of stimulation, ii) if they can accelerate learning and iii) if they would accumulate over several days of stimulation. In the present study, we investigated the lasting behavioural and electrophysiological effects of applying rhTMS over the left intraparietal sulcus (IPS) throughout the course of seven sessions of cognitive training on an auditory working memory task. Fourteen neurologically healthy participants took part in the training protocol with an auditory working memory task while being stimulated with either theta (5Hz) rhTMS or sham TMS. Electroencephalography (EEG) was recorded before, throughout five training sessions and after the end of training to assess to effects of rhTMS on behavioural performance and on oscillatory entrainment of the dorsal auditory network. We show that this combined approach enhances theta oscillatory activity within the fronto-parietal network and causes improvements in auditory working memory performance. We show that compared to individuals who received sham stimulation, cognitive training can be accelerated when combined with optimized rhTMS, and that task performance benefits can outlast the training period by up to 3 days. Furthermore, we show that there is increased theta oscillatory power within the recruited dorsal auditory network during training, and that sustained EEG changes can be observed up to 3 days following stimulation. The present study improves our understanding of the causal dynamic interactions supporting auditory working memory. Our results constitute an important proof of concept for the potential translational impact of non-invasive brain stimulation protocols and provide preliminary data for developing optimized rhTMS and training protocols that could be implemented in clinical populations.

## 1 Introduction

Working memory, the ability to manipulate information stored within short-term memory, is a fundamental cognitive function that underlies verbal comprehension and many other complex processes (Baddeley, 2003; Cowan, 2008). Deficits or decline in working memory are pronounced in normal aging (Belleville et al., 1998; Grady, 2012) as well as in in developmental and degenerative disorders, such as attention deficit hyperactivity disorder (Kofler et al., 2018) and Alzheimer’s disease (Stopford et al., 2012), as well as schizophrenia (Forbes et al., 2009) and human immunodeficiency virus infection (Chang et al., 2001). Working memory is especially important within the auditory domain because the temporally transient nature of sound waves requires the listener to process, maintain, and operate on auditory information in real time (Albouy et al., 2013; Albouy et al., 2015; Albouy et al., 2019; Foster et al., 2013; Kumar et al., 2016; Zatorre et al., 1994).

Several studies have demonstrated that long-range connections between the temporal, parietal, and frontal lobes underly the processing, retention, and manipulation of auditory information (Lewis & Van Essen, 2000; Rauschecker & Scott, 2009). What’s more, there is anatomical and functional evidence to support the dissociation of ventral and dorsal streams within auditory processing, analogous to the dual processing scheme in visual perception (Gallivan & Goodale, 2018; Rauschecker & Tian, 2000). The ventral auditory pathway is defined as relaying object-related information from primary auditory cortex (A1) to anterior temporal and inferior frontal cortex (Petrides, 2014). For example, it has been shown that simple pitch retention recruits the ventral auditory pathway (Albouy et al., 2015; Kumar et al., 2016; Warren & Griffiths, 2003).

By contrast, the dorsal pathway projects sensory information from A1 to posterior parietal cortex and thence to premotor areas (Petrides, 2014) for a variety of cognitive computations, including spatial and sensorimotor transformations (Rauschecker, 2018). The posterior parietal cortex is a key hub for spatial attention and working memory for order (Grefkes & Fink, 2005; Marshuetz et al., 2000; Wager & Smith, 2003). More specifically, the intraparietal sulcus (IPS) has been shown to contribute to manipulating information from multiple sensory domains (Champod & Petrides, 2007; Malhotra et al., 2009; Todd & Marois, 2004; Wendelken et al., 2008). In the auditory domain, these brain regions, in communication with the dorsolateral prefrontal cortex – the frontoparietal network – support the ability to mentally transform auditory information, in the domains of pitch and time (Foster et al., 2013; Schneiders et al., 2012; Zatorre et al., 2010). For example, transposition, that is, modulating the key of a piece of music, can be understood as a type of transformation of the tonal relationships in an abstract pitch space. The IPS within the dorsal pathway has been shown to play an important role for both melodic transposition and melodic reversal (Foster & Zatorre, 2010; Foster et al., 2013).

To understand the role of the frontoparietal network, the neuroimaging literature has historically used correlational approaches to relate patterns of brain activity with behavioral measures. More recent investigations into the causal dynamics supporting cognition have used brain stimulation to modulate the rhythmic firing of large groups of neurons within relevant brain networks (Albouy et al., 2017; Alekseichuk et al., 2016; Di Gregorio et al., 2022; Helfrich et al., 2014; Lakatos et al., 2019; Polania et al., 2018; Riddle et al., 2020; Thut, Schyns, & Gross, 2011; Thut, Veniero, et al., 2011). This work builds upon evidence that neuronal electric field potentials oscillating at specific frequencies can predict an individual’s performance on a cognitive task (Jensen et al., 2007), and that they can be synchronized to an external periodic event (Hanslmayr et al., 2019; Schroeder & Lakatos, 2009; Thut & Miniussi, 2009).

Neural oscillations at different frequencies reflect fundamental elements of brain function: delta (0.5 – 3 Hz), theta (4 – 8 Hz), alpha (8 – 12 Hz), beta (12 – 30 Hz), gamma (30 – 150 Hz) frequency bands are commonly observed in both domain-general and domain-specific cognitive states (Basar et al., 2001; Lopes da Silva, 2013). In addition to frequency, oscillations can be characterized by amplitude and phase, where amplitude depends on excitatory synchrony within local neuronal populations, and phase is related to synchronization of distant brain regions (Fell & Axmacher, 2011). For instance, recent studies have shown that age-related deficits in working memory are associated with reduced theta phase connectivity within the frontoparietal network (Goodman et al., 2019), and that frontoparietal theta desynchronization with noninvasive brain stimulation can induce transient deficits in working memory (Alekseichuk et al., 2017; Polania et al., 2012), Using rhythmic transcranial magnetic stimulation (rhTMS) Albouy et al. (2017) showed that it was possible to modulate these oscillatory patterns during task performance to causally enhance auditory working memory manipulation abilities in healthy adults. rhTMS works by generating localized fluctuations in a magnetic field to modulate the electrical excitability of neurons on the order of milliseconds, via application of trains of repetitive pulses. The authors showed that task-related 5Hz neural oscillatory activity, localized to the IPS, correlated with better performance on a melodic manipulation task resembling those previously shown to engage the frontoparietal network. By administering rhTMS within these functionally relevant parameters – at a frequency of 5Hz and targeted over the IPS – they were able to improve the ability to perform a mental reversal task using brief melodic patterns on a trial-by-trial basis (Albouy et al., 2017). Furthermore, stimulation-associated behavioural benefits (accuracy) were positively correlated with the degree of entrainment resulting from the stimulation. This study therefore established that theta oscillations in the dorsal auditory pathway are causally related to auditory working memory manipulation.

For possible future clinical translation, it is now necessary to show cumulative and long-lasting effects of such procedures, and to understand how non-invasive brain stimulation can interact with cognitive training. Previous efforts to enhance working memory with cognitive training alone have yielded conflicting results, with most training programs producing limited short-term benefits (Malinovitch et al., 2023; Melby-Lervag & Hulme, 2013). Similarly, rhTMS alone has shown promise as an effective intervention for memory improvement, but heterogeneity in stimulation parameters limits its translational potential (Phipps et al., 2021). We propose that a combined protocol of longitudinal cognitive training and rhTMS can produce more durable results if we target the specific oscillatory network dynamics associated with working memory operations. This hypothesis is supported by evidence that repeated stimulation sessions can induce stable late-phase long-term potentiation, a potential mechanism underlying prolonged modulation of network activity and connectivity (Antonenko et al., 2023; Monte-Silva et al., 2013).

There are a growing number of studies showing that transcranial direct current stimulation (tDCS), which uses a constant electrical current to produce sustained modulations of cortical excitability, can also be effectively combined with cognitive training to enhance working memory (Au et al., 2016; Chan et al., 2023; Jones et al., 2020; Nissim et al., 2019; Ruf et al., 2017). Although these studies are promising in terms of their outcomes, by applying tDCS throughout working memory task performance they neglect to consider the oscillatory dynamics of the networks subserving task performance (Andrade-Talavera & Rodriguez-Moreno, 2021; Grover et al., 2021).

Transcranial alternating current stimulation (tACS) offers more precision for cognitive training interventions by using sinusoidal electrical current that can be tuned to the specific frequency of intrinsic brain oscillations (Klink, Passmann, et al., 2020; Klink, Peter, et al., 2020; Lee et al., 2023). With repeated sessions of tACS, Grover et al. (2022) demonstrated long-lasting improvement of auditory-verbal working memory when theta band current was applied to the inferior parietal lobe during encoding and retrieval of word lists. These results encourage our present investigation which applies neuromodulation that is even more tailored to the anatomical location and rhythmic frequency associated with manipulation of information within working memory.

The present study aims to accomplish this goal by administering rhTMS that is optimized to potentiate endogenous oscillations that are specifically and causally related to the training task, an approach described as information-based brain stimulation (Romei et al., 2016). We adapted a working memory training task that can isolate manipulation from simple retention of auditory stimuli (Malinovitch et al., 2023), and we apply rhTMS only during a 2 second time window in between stimulus presentation, informed by dynamic spatiospectral changes during manipulation. Our longitudinal study design allows us to monitor learning over time and persistence of any rhTMS-induced oscillatory and/or behavioural changes.

In addition to testing for behavioural improvements on the trained auditory working memory task, the present study also looked for evidence of near and far cognitive transfers to an auditory working memory task with noise stimuli (near transfer) and to a visual mental rotation task (far transfer). Several functional imaging studies have reported increased activity in the left IPS during visual mental rotation (Alivisatos & Petrides, 1997; Cohen et al., 1996; Hiew et al., 2023; Jordan et al., 2001) and there is strong evidence implicating theta activity within the frontoparietal network in manipulation of both auditory and visual information (Kawasaki et al., 2014). Therefore, it is plausible that strengthening the oscillatory activity within this network via brain stimulation applied during auditory working memory training may induce a far transfer effect to tasks that share the same neural substrates even if they belong to distinct sensory domains. In fact, Albouy et al. (2022) obtained evidence in favor of shared neural resources between visual mental rotation and auditory working memory manipulation by showing that visual stimulation with rotating shapes enhanced performance on a tonal reordering task.

Furthermore, we improve upon previous studies by using image-guided stimulation adjusted to participants’ own cortical anatomy. Our protocol also includes concurrent electroencephalography (EEG) recording before, after, and during 5 training sessions, which addresses a limitation in many prior published studies that do not document the consequences of stimulation on neurophysiological measures. EEG data allow us to measure the immediate after-effects of rhythmic brain stimulation on entrainment of neural oscillations, and thus to document whether the stimulation had the intended effect, and to what extent that was the case. We hypothesize that the combination of information based rhTMS with training will accelerate and augment learning on our auditory working memory task, compared with sham stimulation, and that the changes in behavior will be mirrored in theta-band power on an individual basis.

## 2 Materials and Methods

### 2.1 Participants

We retained 18 neurologically healthy young adults (8 women; mean age of 22 years, ranging from 18 to 29 years) to participate in our study after screening (see below). All participants were right-handed and reported normal hearing and no history of neurological disease. Nine participants had some musical training, with an average duration of 2.5 years and no participant having more than 4 years of training. They gave their written informed consent and received monetary compensation for their participation. Ethical approval was obtained from Ethics Review Board of the Montreal Neurological Institute (NEU-14-043).

Prior to enrollment, eligible participants completed a 10-minute in-person screening test for their ability to perform an auditory working memory task (shown in Figure 1A and described in section 2.3) above chance level. Behavioural data from this screening test were analyzed using the Hits – False Alarm rate. To avoid a potential ceiling effect, an additional exclusion criterion was defined as a Hits – False Alarm rate above 70% on the screening test.

**Figure 1:**
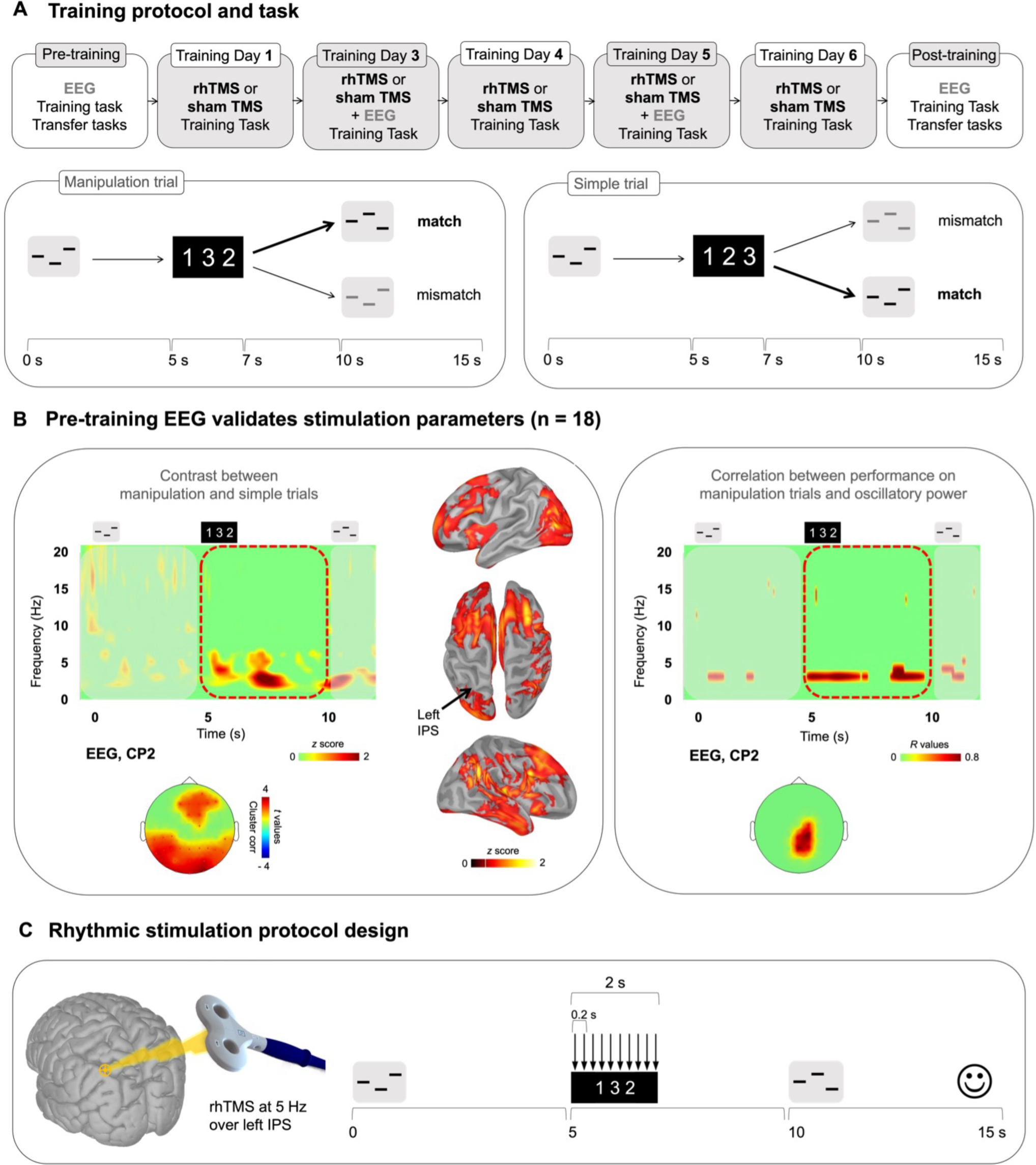
**(A) Top panel:** Overall training protocol. 1 pre-training session, 5 training sessions, and 1 post-training session. All sessions were identical for the 2 groups (rhTMS and sham). **Bottom panel:** Auditory working memory training task. Timeline of a trial in the working memory task as administered in all sessions. Participants listened to a sequence of 3 pure tones. A visual string displaying the expected order of tones in the second sequence was presented 5s after the onset of the first sequence. After 5 additional seconds, participants heard another 3-tone sequence, composed of the same tones, and had to determine whether the order of its tones matches the order of the visual string. Correct responses are bolded for an example of a simple trial (left) and manipulation trial (right) **(B**) Electrophysiological data from the pre-training session. **Left panel:** Time-frequency maps (all participants, n=18) of EEG electrode CP2 for a trial time window (−100 to 12 000ms) for the difference Manipulation versus Simple trials. Red dotted outlines indicate the time period of interest (Manipulation period from 5000ms to 10 000ms post-stimulus onset). Time-frequency maps were *Z* scored with baseline activity (−1 000 to 0ms before stimulus onset). Cortical surface renditions show the difference (z-score) between Manipulation trials compared to Simple trials. Scalp topography show significant clusters where the power of theta oscillations was higher for the Manipulation trials compared to Simple trials for the entire manipulation time window (average over time between 5 and 10s). **Right panel:** EEG topography and time-frequency representation of correlation scores (*R* values) between the power of EEG signals shown at representative electrode CP2, and behavioral performance (d’) for Manipulation trials. **(C)** Stimulation parameters defined by the results presented in (B).

Of 32 participants screened, 19 participants were enrolled for 7 experimental sessions taking place 48-72 hours apart. An overview of the training protocol is depicted in Figure 1A. One participant was terminated after the pre-training session due to below chance performance on simple trials of the working memory task (see below and Figure 1A for task description), yielding a final sample of 18 participants for assignment to experimental and sham conditions as described in section 2.5.

### 2.2 Stimuli

The auditory stimuli (taken from Malinovitch et al. (2023)) used in the working memory training task consisted of sequences of three 400 millisecond pure tones randomly selected from a frequency range of 250 to 1000 Hz, with tones in the same sequence having frequencies at least 20% different from one another. These sequences were delivered binaurally through air-conducting tubes with foam ear tips (70 dB SPL).

### 2.3 Training task

The auditory working memory task (from Malinovitch et al. (2023)) administered during the screening test and throughout the 7 experimental sessions involved the mental manipulation of the order of three pure tones and is depicted in Figure 1A. In each 15-second trial, a sequence of three tones was presented, followed by a silent retention period of 3800 milliseconds. A visual cue then appeared on the screen, consisting of the numbers 1, 2, and 3 in variable order. The order of these numbers corresponded to the order of tones in a reordered target sequence. For example, the visual cue “312” instructed participants to reorder the sequence of the original tones (say, A-B-C) such that the third tone now came first, that is, C-A-B. Finally, after 5 seconds participants were presented with a second tone sequence and indicated by left or right click on a mouse whether it was a ‘match’ or ‘mismatch’ to the target sequence, that is, the correct manipulation of the encoded sequence. At the end of each trial, visual feedback indicated whether the response was correct.

An important aspect of the task design was the intermixing of two different types of trials: simple and manipulation trials. A simple trial is any trial wherein the visual instruction is “123”. In such trials, participants are not required to perform any mental reordering of musical tones; they must simply judge whether the second sequence is the same as or different from the encoded sequence. This melodic comparison does not involve working memory, so it can be classified as a short-term memory task. On manipulation trials, the visual instructions are not in numerical order, such as “213” or “321”, indicating that participants must perform a manipulation on the auditory information stored within short-term memory. The contrast between these two trial types allows us to identify neurophysiological properties that are unique to manipulation abilities.

There were 42 consecutive trials (14 simple and 28 manipulation) within one 10-minute run, and three runs within each 30-minute testing session for a total of 126 trials. Within each run, the trials were presented in a fixed randomized order. The runs presented in the pre-training and post-training sessions were identical, while unique runs were generated for each training session (but were similar for all participants). The order of the runs in each session was counterbalanced across participants.

### 2.4 Transfer tasks

On pre-training and post-training sessions we administered a working memory task with identical design to the training task, except with different auditory stimuli, to assess evidence of near transfer. The stimuli used in this transfer task, which we refer to as the noise task (Supplementary Figure 1A) were comprised of 42 different sequences of three unfamiliar non vocal sounds, or “noises”. Each noise sequence was comprised of 400 millisecond audio clips of similar overall energy (RMS) that had no recognizable tone (material from Malinovitch et al. (2023), 50% match trials and 50% mismatch trials). Note that there were only manipulation trials in this near transfer task.

An additional visual mental rotation task was administered after the auditory working memory task, exclusively in experimental sessions 1 and 7 to test for far cognitive transfer. Two three-dimensional asymmetrical forms, each comprised of 7-9 cubes as in Shepard and Metzler (1971) were displayed side by side on a computer screen. The 10 visual stimuli used in this task – three 7-cube shapes, six 8-cube shapes, and one 9-cube shape – were composed of white cubes on a black background (see Supplemental Figure 1B). The paired shapes were either identical or the left-hand shape was horizontally mirrored. Relative to the shape on the left, the right-hand shape was rotated around its own axis by either 0, 60, 120, 180, 240, or 300 degrees.

Each trial began with the presentation of a fixation cross for 250 milliseconds, followed by the paired target image and rotated probe. Participants were instructed to indicate, using left or right arrow keys, whether the two forms were identical or mirror images of one another after mental rotation. No feedback was delivered, and subsequent trials occurred immediately after a response was recorded, or after a delay period of 8 seconds if no response was recorded. The task began with a practice phase comprising 10 trials for which responses were not recorded, followed by the experimental data collection phase: all 10 stimuli pairs were presented once at each of the six relative rotation angles for both mirror and non-mirror trials, for a total of 120 trials in random order. One full run of the task took a maximum 16 minutes to complete.

### 2.5 Experimental protocol

#### 2.5.1 Anatomical data

All participants underwent a 3D anatomical MPRAGE T1-weighted Magnetic Resonance Imaging (MRI) on a 1.5T Siemens Sonata scanner or on a 3T Siemens Trio (Siemens AG, Munich, Germany) before or just after the EEG recording of the pre-training session. Diffusion weighted imaging (DWI, data not presented here) was also acquired during this session. The anatomical volume consisted of 160 sagittal slices with 1 mm^3^ voxels, covering the whole brain. The scalp and cortical surfaces were extracted from the T1-weighted anatomical MRI. A surface triangulation was obtained for each envelope using the segmentation pipeline available in FreeSurfer (Fischl, 2012) with default parameter settings. The individual high-resolution cortical surfaces (about 75 000 vertices per surface) were down-sampled to 15 002 vertices using Brainstorm (Tadel et al., 2011) to serve as image supports for EEG source imaging.

#### 2.5.2 EEG recording

For all recordings we used TMS-compatible EEG equipment (two 32 channel BrainAmp DC amplifiers, BrainProducts, http://www.brainproducts.com/). EEG was continuously acquired from 62 channels (plus ground, EOG and nose reference electrodes). TMS-compatible sintered Ag/AgCl-pin electrodes were used. The signal was band-pass filtered at DC to 1 000 Hz and digitized at a sampling rate of 1 000 Hz. Skin/electrode impedance was maintained below 10 kΩ. The positions of the EEG electrodes were estimated using the same 3D digitizer system (Polhemus Isotrack). Most EEG pre-processing, EEG source imaging and statistical analyses were performed with Brainstorm (Tadel et al. (2011), http://neuroimage.usc.edu/brainstorm/) combined with Fieldtrip functions (http://www.fieldtriptoolbox.org/) and custom-made MATLAB code.

#### 2.5.3 TMS Protocol

On training days (i.e. experimental sessions 2 to 5), TMS was applied during task performance and during EEG recording. Participants were seated with their chin positioned in a chin rest, their eyes open, and their gaze centered on a continuously displayed fixation cross (white on a gray background). They listened to the auditory stimuli presented binaurally through air-conducting tubes with foam ear tips (70 dB SPL). They were asked to maintain central fixation and to minimize eye blinks and other movements during the recording blocks. Short biphasic TMS pulses were delivered by a TMS coil (70mm figure-eight coil connected to a Magstim Rapid2 Stimulator) over the left Intraparietal Sulcus for the rhTMS group or was tilted 90° away from the head for sham-TMS group (control group) For the rhTMS group, the TMS coil was oriented perpendicular to the target region, to maximize effect strength (Thut, Veniero, et al., 2011). We identified the IPS (marked at x −36 y - 60 z 56 in MNI space, coordinate from Albouy et al. (2017)) using neuronavigation (Brainsight, Rogue Research Inc, Montreal, Canada) based on the participant’s own MRI scan in the native space. The transformation of the MNI coordinate to the native space was done using SPM 12 normalization functions.

For each trial, for both groups, ten TMS pulses were delivered during the manipulation period (onset of the visual instruction, i.e. 5 s after the onset of the first sound sequence). The TMS pulses were delivered at 5Hz (1 pulse every 200ms, frequency of stimulation defined with EEG data of the pre-training session, see Results, Figure 1B, C). TMS intensity was at 60% of machine output (see Albouy et al. (2017); Weisz et al. (2014) for similar procedure and Thut, Veniero, et al. (2011) with TMS intensity ranging from 58% to 66%).

On each TMS session there were three blocks of the task. In each block, 42 ten-pulse TMS trains were delivered, leading to 420 pulses per block over a block duration of about 10 min. Each TMS/EEG session (sessions 2 to 5) thus contained a total of 2160 active TMS pulses. The frequency of the rhythmic TMS was fixed to 5Hz (based on EEG results), and therefore was not adjusted to each participant’s individual theta frequency. This stimulation parameter was chosen based on previous rhythmic TMS studies showing that individual frequency tuning may not be a strict requirement for entrainment (see Romei et al. (2010)). Notably, it has been shown that with increasing stimulus intensity, the relationship between the effective stimulation frequency and the preferred frequency tends to be reduced (Glass, 2001). With such strong driving forces as TMS, entrainment may be enabled using a relatively large frequency range (see Thut, Veniero, et al. (2011)), and therefore it may not be required to set the TMS rate to each participant’s individual self-generated frequency to observe behavioral effects. The TMS protocol respected the safety recommendations regarding stimulation parameters (intensity, number of pulses, ethic requirements) presented in Rossi et al. (2009).

#### 2.5.4 Procedure

##### Pre-training session

Participants performed the working memory task for tones, the working memory task for noises and the mental rotation task in the absence of stimulation, and with EEG signal recording. For the working memory task for tones, we computed the Hits – False Alarm rate for the manipulation trials within the three runs, as a measurement of baseline performance (i.e., performance at the pre-training session). Participants were then assigned into either 5Hz rhTMS or sham rhTMS groups, balancing for gender and baseline task performance.

##### Training days

Participants completed three runs of the auditory working memory task while receiving real or sham rhTMS during training days 1-5 (Fig 1A). On training days 2 and 4 we additionally obtained concurrent EEG recordings to assess the impact of rhTMS on oscillatory entrainment of the neural network engaged in task performance.

##### Post-training session

During the post-training experimental session, participants again performed the working memory task for tones, the working memory task for noises, and the mental rotation task in the absence of stimulation, and with EEG signal recording. We administered an exit survey for all participants upon finishing the experiment. Participants were asked to report their perception of how much they improved throughout the course of training and to describe any cognitive strategies they employed to perform the auditory manipulations, such as visualization or counting the position of the highest tone in the sequence.

### 2.6 Data analysis

#### 2.6.1 Behavioural data

Behavioural data were analyzed using signal detection theory in order to measure discrimination ability between correct and incorrect manipulations, unbiased by tendency to respond “match” or “mismatch” more frequently. For this analysis, a hit is considered a “mismatch” response to a mismatch trial, and a false alarm is a “mismatch” response to a “match” trial. We computed d’ values to compare the average performance on manipulation trials between experimental groups and across days. Note that the same analysis strategy was used for the simple task.

#### 2.6.2 EEG pre-processing and TMS artifact removal

Following procedures previously described in Albouy et al. (2017), preprocessing was performed in multiple steps, starting with removal of bad segments by visual inspection, and removal of the dominant TMS artifacts for EEG data of sessions 3 and 5. For this purpose, TMS artifacts were automatically detected and a period starting 10ms prior to and ending 20ms after the respective TMS peak was replaced by Gaussian noise with the standard deviation and mean adapted to correspond to a reference period set to be −35 to −15ms before the respective TMS peak (see Albouy et al. (2017)). Following this step, the data were down sampled to 500 Hz. This procedure effectively removes the direct (non-physiological) TMS artifact without introducing discontinuities, important for the later time–frequency analysis (see Weisz et al. (2014), Thut, Veniero, et al. (2011)). However, this measure still left some TMS locked artifacts at electrodes directly in contact with the TMS coil.

These residual artifacts were effectively removed using Independent Component Analysis (ICA) using EEG lab functions (https://sccn.ucsd.edu/eeglab/). For this purpose, as well as for the removal of artifacts of other origin (eye movements/blinks) artifact rejection removing was ran (e.g., dead channels, channel jumps, etc.) and the data was subsequently filtered between 0.3 to 50 Hz before computing the ICA on the remaining data. Using time-course and topographic information, components representing clear ocular or TMS-related artifacts were identified and removed from the filtered data. In a last preprocessing step, residual artifactual trials were removed by visual inspection. Note that the TMS artifact correction procedure has been evaluated in a control experiment (see Figure S5 in Albouy et al. (2017). EEG data of sessions 1 and 7 were pre-preprocessed with a similar procedure, but without correcting for TMS artifacts. The files were re-referenced to the average of all channels. Individual EEG trials were then automatically inspected from −1 000ms to 12 000ms with respect to the onset of the first tone of the first sequence. Trials with ranges of values exceeding ± 250 μV within a trial time-window at any electrode site were excluded from the analysis.

#### 2.6.3 EEG source imaging

Source reconstruction was performed using functions available in Brainstorm, all with default parameter settings (Tadel et al., 2011), as in Albouy et al. (2022). Forward modeling of neural magnetic fields was performed using a realistic head model: symmetric boundary element method from the open-source software OpenMEEG. A realistic BEM model of head tissues and geometry for the anatomy of each participant (see Anatomical data) was used, as EEG data are sensitive to variations in head shape and tissue conductivity. The lead fields were computed from elementary current dipoles distributed perpendicularly to the cortical surface from each individual. EEG source imaging was performed by linearly applying Brainstorm’s weighted minimum norm operator onto the preprocessed data. The data were previously projected away from the spatial components of artefact contaminants. For consistency between the projected data and the model of their generation by cortical sources, the forward operator was projected away from the same contaminants using the same projector as for the EEG data. The EEG data were projected on a cortical surface in the native space (cortical surface of 15 002 vertices serving as image support for EEG source imaging).

#### 2.6.4 Oscillatory activity

We were first interested in confirming the role of theta oscillations during the manipulation of information in memory (as compared to simple retention). We thus focused on theta activity during the manipulation period between the presentation of the visual instruction and the onset of the second sound sequence. We performed wavelet time-frequency decompositions of sensor signals (Tallon-Baudry et al., 1996). The EEG signals were convoluted with complex Morlet’s wavelets, with a Gaussian shape in both the time (SD σ*t*) and frequency domains (SD σ*f*) around their central frequency *f*0. The wavelet family was defined by (*f*0/σ*f*) = 7, with *f*0 ranging between 1 and 80 Hz in 1 Hz steps. The time-frequency wavelet transform was applied to each trial and then averaged across trials, resulting in an estimate of oscillatory magnitude at each time sample and at each frequency bin between 1 and 80 Hz. Time-frequency decompositions of signal during the period were *z*-scored with respect to a pre-stimulus baseline (−1 000 to 0ms before the presentation of the first tone of the first sequence).

For session 1, with 18 participants, the resulting time-frequency maps were correlated to the individual behavioral performances (correlation applied at each frequency band and time sample), as illustrated in Figure 1B. For all sessions, EEG signals were filtered in the theta frequency band (4 to 8 Hz) before their envelope was extracted using the Hilbert transform. The resulting signal magnitude envelopes were baseline-corrected using *z* scores with respect to the mean theta power over −1 000 to 0ms preceding the presentation of the first tone of the first sequence. For each session and each participant, we derived an averaged version of these data for the period between 5 000 to 10 000ms corresponding to the manipulation period. The resulting maps were then contrasted (manipulation vs. simple for session 1, group comparison for sessions 2 to 7, and post vs. pre contrast for each group). Statistical significance was tested using cluster-level statistics (alpha = 0.05, one tailed see Oostenveld et al. (2011). We then performed the same analyses (Hilbert, average 5-10s, cluster permutation testing) at the source level.

## 3 Results

Of the 18 participants included in our dataset, 14 completed all 7 experimental sessions. The remaining 4 participants had partially completed the training protocol when data collection was interrupted by the SARS-CoV-2 pandemic. Accordingly, 14 participants are included in the comparative group analyses investigating the online (during stimulation) and lasting effects of rhTMS combined with cognitive training. Thus, the following results compare both auditory working memory performance and neural oscillatory dynamics within the task-recruited network, between stimulation and sham control groups, with 7 participants in each rhTMS and sham control groups.

### 3.1 Theta activity in the dorsal pathway is greater during manipulation compared to simple trials and is positively correlated with pre-training performance

We used EEG data from the entire sample of 18 participants who completed the pre-training session to validate that parietal theta activity was associated with manipulation abilities in our dataset. Time-frequency maps were generated for correct manipulation trials and then baseline corrected and averaged over trials for each participant (see methods). The same procedure was applied to correct simple trials. We then subtracted the averaged simple trials from the averaged manipulation trials for illustration, to isolate the activity elicited by mentally reordering the auditory stimuli and averaged the difference maps across all participants. In the resulting averaged difference map shown in Figure 1B, a burst of theta band activity coincided with presentation of the visual cue and was sustained until presentation of the test auditory stimuli. The corresponding scalp topography and source reconstruction for the manipulation time period (5-10 seconds) show prominent theta power (4-8Hz) over the posterior parietal cortex and frontal cortices. Note that this difference was significant as illustrated in the scalp topography in Figure 1B (cluster corrected, alpha = 0.05, frontal cluster, k = 7, p = .01, parietal cluster, p = .004, k =23).

Additionally, individual participants’ theta oscillatory power during this time period of 5-10 seconds was positively correlated with accuracy on manipulation trials (statistical peak r(17) = .85, p< .001). As shown in Figure 1B, we generated a time-frequency map of the correlation between oscillatory activity, averaged across pre-training manipulation trials, and d’ on pre-training manipulation trials. Highly behaviorally correlated theta (4-8Hz) power during the manipulation period (5-10s) was concentrated in parietal electrodes.

### 3.2 Rhythmic TMS increases the rate of improvement during working memory training

A two-tailed Mann-Whitney U-test indicated that rhTMS and sham control groups did not differ in terms of the discriminability index (d’) on manipulation trials at pre-training (U = 21, p = 0.71). Similarly, there was no group difference when looking at simple trials (U = 11, p = 0.1). We plotted the average d’ of both groups in each experimental session (Figure 2). To compare the rate of learning between groups, we generated a simple linear fit (linear regression) of each participant’s accuracy (d’) in training sessions 2-6. We then extracted the slope of the linear fit for each participant (Figure 2), reflecting the rate of change in accuracy over sessions and conducted a two-tailed Mann-Whitney U-test that revealed a significant group difference for manipulation trials (U = 9, p = 0.05) but not simple trials (U = 16.5, p = 0.34).

**Figure 2:**
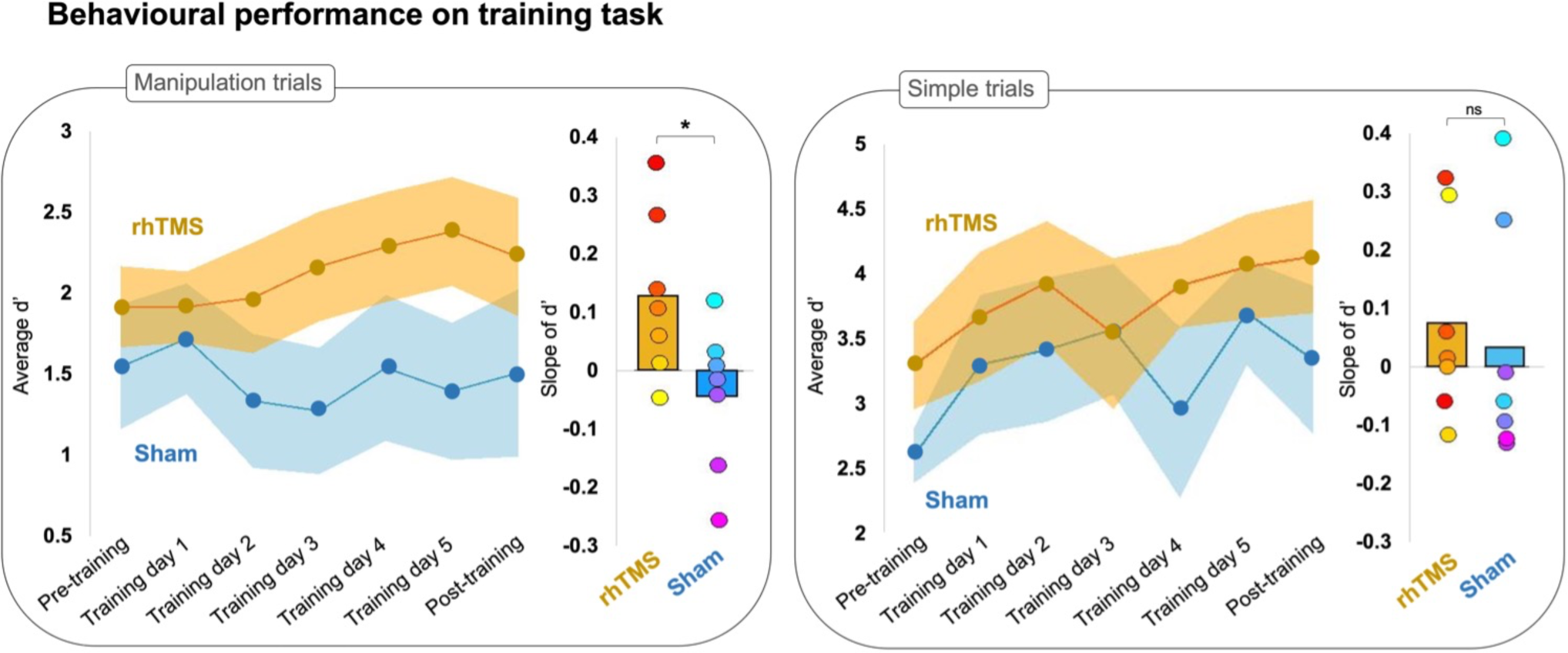
Line graphs plotting average accuracy (d’) on manipulation (left) and simple (right) trials of an auditory working memory task over 7 experimental sessions for rhythmic stimulation (rhTMS, n=7, orange) and control (Sham, n=7, blue) groups. Shading represents SEM. Bar plots of the average slope extracted from a simple linear fit for individual d’ data points from manipulation (left) and simple (right) trials on training days, by experimental group. Asterisks indicate significance.

We further investigated this finding with a one sample Wilcoxon signed rank test and found that participants stimulated with rhTMS had on average a manipulation trials learning slope significantly greater than zero (W = 26, p = 0.02), whereas participants in the sham condition had on average a slope of learning on manipulation trials that was not greater than zero (W = 9, p = 0.81). Neither experimental group had a slope greater than zero for the d’ on simple trials (rhTMS W = 14.5, p = 0.23; sham W = 13, p = 0.59). We then performed a one-tailed Wilcoxon signed rank test of d’ at pre-training vs. post-training, which showed that the rhTMS group exhibited a significant post-training increase in performance on manipulation trials as compared to pre-training (W = 4, p = 0.05), but that the sham control group did not have significantly improved performance on manipulation trials at post-training compared to pre-training (W = 16, p = 0.66). The same analysis applied to performance on simple trials revealed a similar pattern, with the rhTMS group exhibiting an improvement at post-training (W = 0, p = 0.01) but not the sham group (W = 5, p = 0.08). Note however, that the contrast rhTMS group vs. sham group was not significant for post-training manipulation trials (U = 19, p = 0.54) or simple trials (U = 16.5, p = 0.33). We did not find any group differences in behavioural performance on either the noise or mental rotation tasks. The data and statistics for each task are reported in Supplemental Figure 1.

### 3.3 Rhythmic TMS elicits greater oscillatory entrainment compared to sham TMS during training

We next compared task-relevant theta oscillatory activity between groups during the two training days with concurrent EEG recording (training days 2 and 4) because we were interested in evaluating the oscillatory entrainment in participants receiving real and sham rhTMS. Time-frequency maps were generated for correct manipulation trials, baseline corrected, then averaged across both training days for each participant. With these individual maps we constructed averaged time-frequency maps for both the rhTMS group and the sham group (Figure 3A).

**Figure 3:**
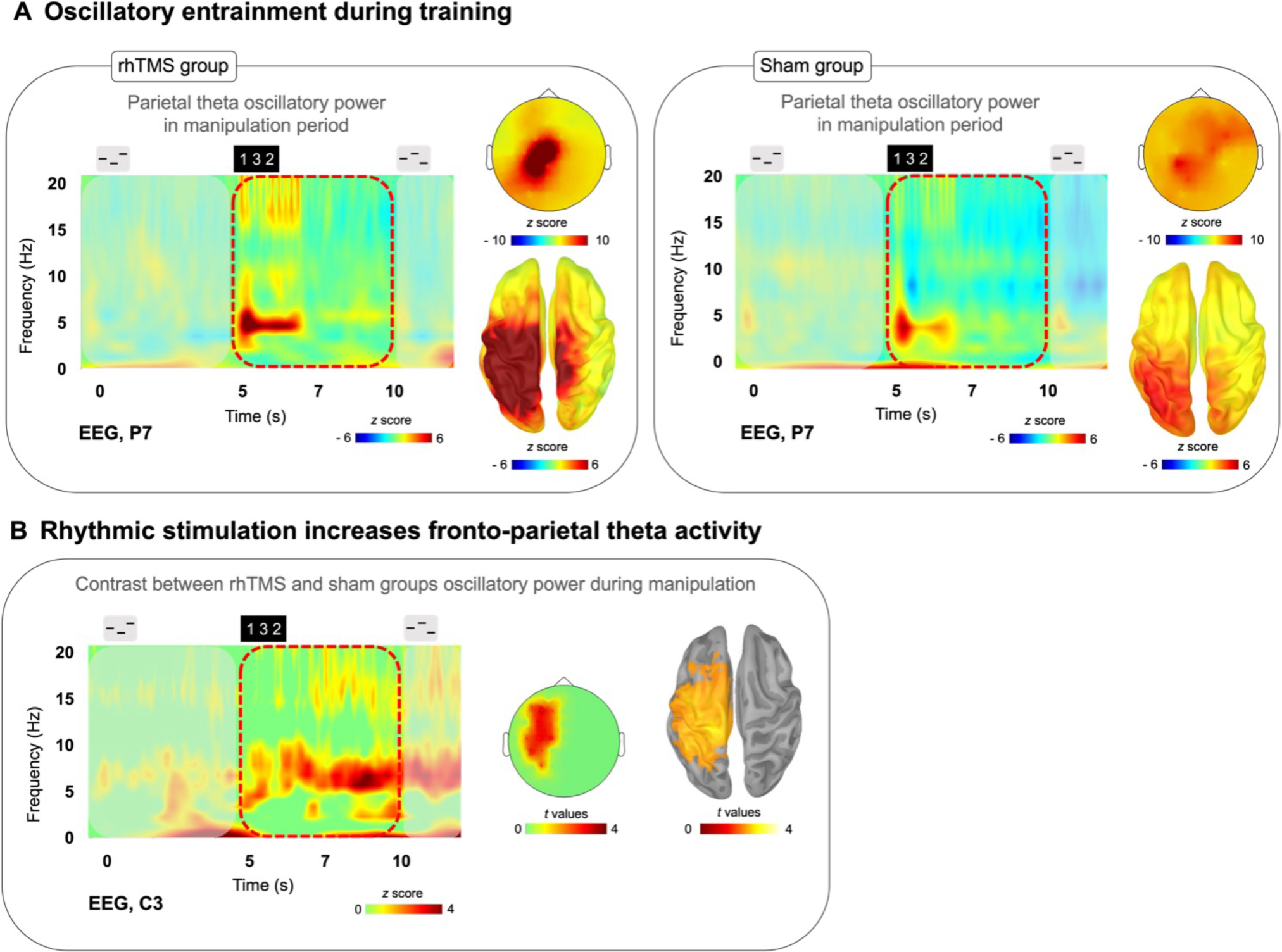
**(A)** Time-frequency plots (P7 electrode) for a trial time window with scalp topography and source localization from the average of training days 2 and 4 manipulation trials for both rhTMS (n=7, left panel) and Sham (n=7, right panel) groups, showing sustained theta band (4-8Hz) activity in parietal regions from 5-10s in both groups. **(B)** Time-frequency plots with statistical tests performed at the sensor and source levels for the contrast rhTMS vs. sham group, showing significant clusters of theta band activity in fronto-parietal regions for the average of the entire manipulation time period (5-10s) of manipulation trials (averaged across training days 2 and 4).

To evaluate group differences in theta oscillatory power within the manipulation period of auditory working memory task performance, we conducted a one-tailed t-test with cluster correction for the difference in the theta (defined as the 4-8Hz frequency band) oscillatory activity, over the time window of 5-10 seconds in correct manipulation trials from sessions 3 and 5, between rhTMS and sham groups. A significant cluster of theta activity was revealed at the sensor (cluster corrected, alpha = 0.05, k = 8, Figure 3B) and source levels (cluster corrected, alpha = 0.05, p = 0.05, k = 3308 vertices, Figure 3B) overlying the left fronto-parietal network.

### 3.4 Post-training oscillatory entrainment is greater following repeated sessions of rhTMS compared to sham TMS

Next, we tested our main hypothesis that combined working memory training and rhTMS would induce a lasting enhancement of task-relevant oscillatory dynamics, specifically theta oscillatory power in the fronto-parietal network. For illustration, we performed a simple subtraction of individual pre-training manipulation trial time-frequency maps from the corresponding post-training maps; the resulting contrast maps for each group are displayed in Figure 4A.

**Figure 4:**
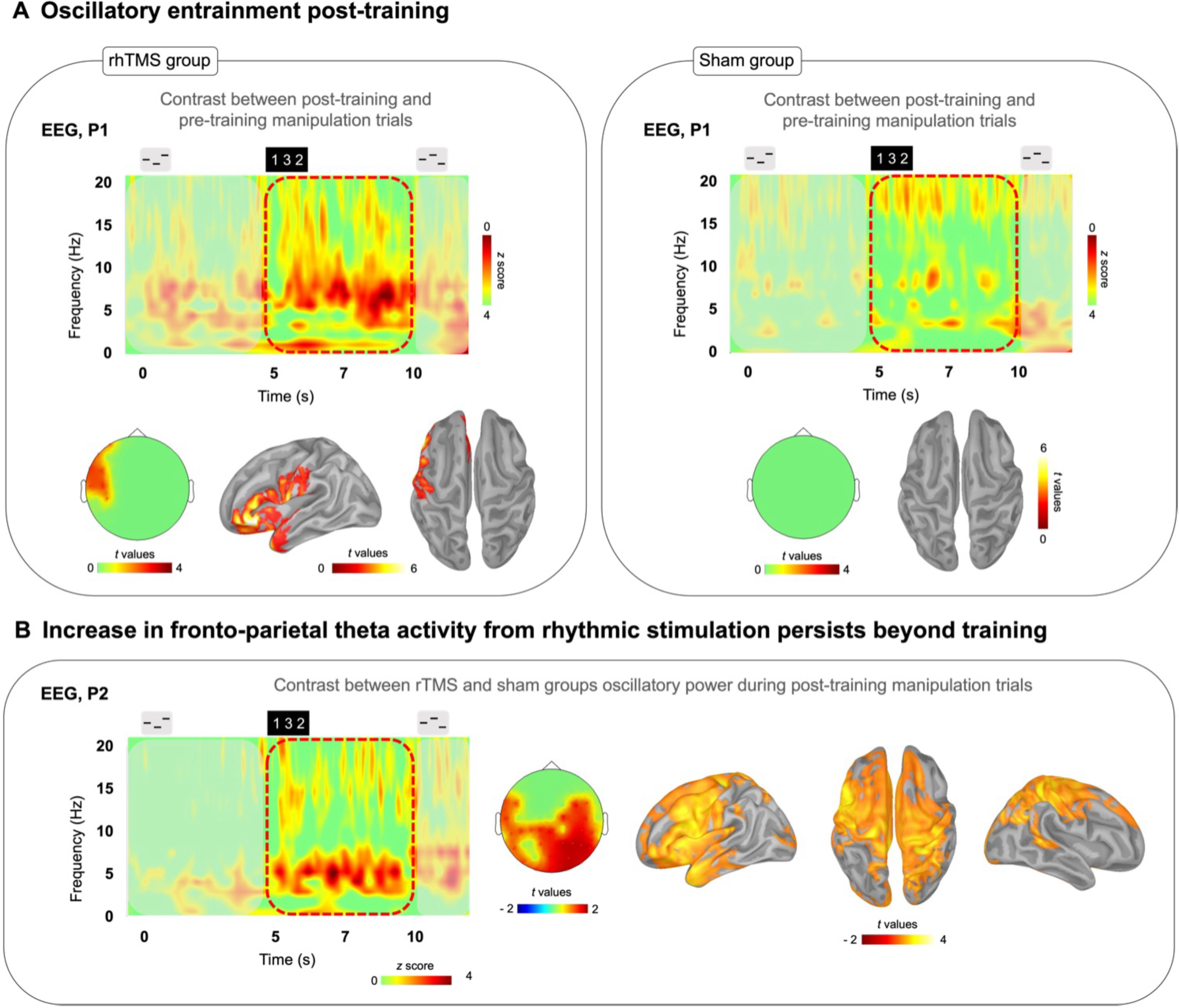
**(A)** Time-frequency plots of the difference of post-training correct manipulation trials minus pre-training correct manipulation trials for both rhTMS (n=7, left panel) and Sham (n=7, right panel) groups. **Left**: Significant cluster of left frontal theta band (4-8Hz) activity from 5-10s in the rhTMS group for the contrast post vs. pre-training. **Right**: No significant differences in theta band activity between pre and post-training in the sham group. **(B)** Time-frequency plot of the difference resulting from subtracting sham group from rhTMS group post-training manipulation trials. Scalp topography and source localization showing significant group differences in theta band activity in fronto-parietal regions throughout the entire manipulation time period (5-10s) of manipulation trials post-training.

We then measured the change in theta oscillatory activity by conducting a paired one-tailed t-test with cluster correction contrasting theta (defined as the 4-8Hz frequency band) oscillatory activity, over the time period of 5-10 seconds in correct manipulation trials, between post-training and pre-training sessions. A significant cluster of theta activity was revealed in left frontal cortex at the sensor (cluster corrected, alpha = 0.05, p = 0.01, k = 7) and source level (cluster corrected, alpha = 0.05, p = 0.035, k = 1898) for the rhTMS group (Figure 4A, left panel), whereas no significant clusters of theta activity appeared for the sham group (Figure 4A, right panel).

Finally, we looked for group differences in oscillatory activity during task performance in the post-training session. Figure 4B shows the time-frequency map produced by subtracting the sham group’s post-training manipulation activity from the rhTMS groups. An independent samples one-tailed t-test with cluster correction for the difference in post-training theta (4-8Hz) activity from 5-10 seconds in correct manipulation trials, between rhTMS and sham groups was significant for clusters overlying distributed frontoparietal regions (cluster corrected, alpha = 0.05, p = 0.01, k = 30, Figure 4B).

## 4 Discussion

### 4.1 Theta power in the dorsal pathway is greater during manipulation compared to simple trials and is positively correlated with pre-training performance

We first confirmed (Figure 1) that frontoparietal theta activity was indeed a useful marker of working memory function in our training task and participant pool by showing that it is specific to the manipulation condition and is positively correlated with individual manipulation abilities. These data confirm and replicate previous findings that implicate the IPS in auditory manipulation (Foster & Zatorre, 2010; Foster et al., 2013) and fronto-parietal theta oscillations in working memory (Albouy et al., 2018; Albouy et al., 2022; Albouy et al., 2017; Fell & Axmacher, 2011; Lisman, 2010; Sauseng et al., 2010). Importantly, in each trial of our working memory task we used a visual instruction at 5s to mark the precise time-point at which participants began to perform the mental manipulation of auditory stimuli, and applied rhTMS accordingly during the trial time window from 5-7s, wherein the dorsal pathway is selectively engaged by endogenously generated oscillatory activity within the theta frequency band. This element of training task design is critical to our central hypothesis, that repeated sessions of rhythmic stimulation, during specific moments in task performance when the relevant networks are engaged, will produce cumulative and durable improvements in network function.

The majority of studies that have combined non-invasive brain stimulation and cognitive training with a similar gain-of-function aim to improve working memory, have used tDCS or tACS methods that are not tailored to endogenous brain rhythms evoked during the training tasks (see Polania et al. (2018) for review). It is difficult to compare these approaches to our findings given the fundamentally different mechanisms of TMS, but one idea that has emerged from the literature is that online brain stimulation, that is, stimulation applied during task performance, is more effective at enhancing or disrupting brain activity than passive brain stimulation protocols (Luber & Lisanby, 2014). We decided that there was minimal value in testing this idea in the present study with an additional control groups receiving rhTMS prior to task performance, and without cognitive training, given the evidence against efficacy of offline 5Hz TMS in healthy adults (Beynel et al., 2019).

### 4.2 Rhythmic TMS increases the rate of improvement during working memory training

First, we observed a significant difference in learning slopes between experimental and sham groups during training, indicating that working memory training can be accelerated with information-based rhTMS (Figure 2). Additionally, we observed a significant behavioural improvement on the trained task between post-training and pre-training sessions in the rhTMS group but not in the sham group (Figure 2). These results suggest that the known performance-enhancing benefits of information-based rhTMS on auditory working memory can persist beyond the period of active stimulation when combined with cognitive training (Alekseichuk et al., 2016; Hanslmayr et al., 2019; Polania et al., 2018; Violante et al., 2017). Their specific novelty is in suggesting that those behavioural improvements are cumulative with repeated sessions and can accelerate learning in a complex auditory working memory task.

The slope of learning during cognitive training was different between groups for manipulation trials. Whereas the rhTMS group had a positive slope, the sham group’s slope was not significantly different from zero. This result was expected and can be explained by making a parallel with the results of Malinovitch et al. (2023) with the same task. In a sample of 28 young healthy individuals, the authors administered an adaptive cognitive training protocol consisting of 40 sessions of training on the same auditory working memory task used in the present study, only with the number of tones in each sequence increasing as participants accuracy increased, beginning with three tones and up to a maximum of 8 tones per sequence. They found that no participants exhibited an improvement on reordering three-tone sequences after the first 5 sessions. Additionally, as the number of tones in a sequence increased, the error rate decreased due to alternative strategies being employed that reduced the cognitive demand for true manipulations within auditory working memory. These findings were used to inform the design of our training task and training protocol, such that we chose to use only sequences of three tones and to limit the number of training sessions to 5, in order to test our hypothesis that information-based rhythmic stimulation would increase the rate of learning.

For learning on simple trials, neither group had a slope that differed significantly from zero. This result is also in line with the previous findings of Albouy et al. (2017) insofar as 5Hz rhTMS specifically enhances performance on manipulation but not simple trials. However, the most likely explanation of the absence of learning on simple trials is related to ceiling effects. It is generally accepted that a range of d’ values between 0.5 and 2.5 is desirable for avoiding obscured results from either floor or ceiling effects, and these sensitivity values roughly correspond to an accuracy rate of 60% and 90%, respectively (MacMillan & Creelman, 2005). The average d’ of sham and rhTMS groups on manipulation trials at baseline was 1.53 and 1.91 respectively, whereas the average d’ on simple trials was higher at 2.61 for sham and 3.3 for rhTMS groups. With such high performance in both groups, there is little room for meaningful improvement.

Unexpectedly, the benefits of rhTMS to behavioural performance did not appear to transfer to two untrained cognitive tasks. At post-training, the rhTMS group did not perform better than the sham group on either the auditory working memory task with noises, or the visual mental rotation task (Supplementary Figure 1). After five sessions of training on the main task, neither experimental group displayed improvement on the untrained noise task. Given that the design of the noise task was identical to the main task apart from the stimuli used, we expected to observe some near transfer of learning, specifically for manipulation of order within the auditory domain. We did however find some evidence of what may be construed as far cognitive transfer to the visual mental rotation task from the auditory task, albeit not as a function of brain stimulation, since both the rhTMS and sham groups displayed significantly improved performance at post-training compared to pre-training, even though they did not practice the visual task. These results could be related to the inherent difficulty of the two tasks; it may require more sessions of cognitive training to see transfer to the noise task, or to see a group difference in transfer to the rotation task. Individual variability in frontoparietal network connectivity could also explain differences in learning generalization to untrained tasks, as in Wards et al. (2023).

### 4.3 Rhythmic TMS elicits greater oscillatory entrainment compared to sham TMS during training

By performing simultaneous rhTMS and EEG recording, we were able to characterize the immediate consequences of rhythmic and sham TMS on oscillatory activity. As shown in Figure 3, and as expected, participants receiving real 5Hz rhTMS had significantly greater entrainment of theta activity in the dorsal pathway than did participants in the sham TMS group. Considering these findings, we believe that our choice to fix the frequency of rhythmic stimulation at 5Hz, unadjusted to each participant’s individual theta frequency, was justified based on previous literature (Albouy et al., 2017). Indeed, our results demonstrate that it is not required to set the rhTMS rate to each participant’s individual self-generated frequency in order to enable entrainment (Figure 3A) and observe behavioral effects (Figure 2).

Subsequent analyses on larger datasets could investigate potential behavioural correlates of specific theta band frequencies generated during task performance. There is currently a lack of evidence supporting different functional significance of high versus low theta band activity. Overall, these effects observed on theta power are consistent with previous studies showing upregulation of oscillatory activity at the target frequency (Romei et al., 2016; Thut, Veniero, et al., 2011; Weisz et al., 2014). The most interesting contribution of the present study to the oscillatory entrainment literature is the investigation of after effects that are discussed below.

### 4.4 Post-training oscillatory entrainment is greater following repeated sessions of rhythmic TMS compared to sham TMS

The results displayed in Figure 4A contrasting theta oscillatory activity in the dorsal pathway during manipulation post-vs pre-training are an exciting reflection of the behavioural changes, with significant differences over time observed only in the rhTMS group. The EEG analysis revealed a significant increase in the 4-8Hz frequency band in the post-training compared to pre-training recordings only in participants who received real 5Hz rhTMS (Figure 4A). Furthermore, post-training theta-band activity was significantly different between the two groups in brain areas corresponding to the frontoparietal network (Fig 4B). Taken together, we can say that training with rhTMS caused an increase in theta power that outlasted the period of stimulation by 48-72 hours. This evidence implicates neuroplastic mechanisms, specifically induced by rhTMS, underlying the prolonged up-regulation of oscillatory activity within the dorsal pathway.

Previous efforts to explore the relationship between non-invasive brain stimulation, neuroplasticity, and the enduring impact on neural circuitry, have largely focused on Hebbian mechanisms like long-term potentiation and long-term depression (Cirillo et al., 2017). There remains a lack of direct verification for the physiological consequences of TMS protocols for enhancing cognitive function in healthy subjects. While the complex cellular and molecular mechanisms of plasticity that are modulated by TMS, among other methods of noninvasive brain stimulation, are presently best elucidated by *in vivo* and *ex vivo* animal studies, we can obtain indirect, systems level evidence from human clinical studies, albeit with notable inter and intra individual variability (Jannati et al., 2023). This multidisciplinary research supports our finding that precisely timed pulses of magnetic stimulation and subsequent entrainment of neural rhythms can be combined with cognitive training to facilitate consolidation of learning in specific neural networks.

Drawing upon the literature, one possible explanation for how this might occur is that during training, the ongoing intrinsic theta rhythms become phase-locked to exogenous rhythmic stimulation and that this amplifies signals of cortical synaptic plasticity. By virtue of the strong resonant frequency, theta oscillatory activity is consistently boosted in frontoparietal brain regions during task performance. Over repeated trials and sessions, the enhanced synchronous cortical activations reinforce communication between frontal and parietal areas. Long-lasting changes to tissue microstructure thereby improve the efficiency of information flow along relevant white matter pathways, a process that would organically occur much more slowly in the absence of rhTMS.

We have shown here that cognitive training combined with rhTMS accelerates learning and potentiates endogenous theta oscillatory activity within the frontoparietal network. In healthy individuals tested 48-72 hours after receiving a 5-day cognitive training program with information-based rhTMS, there is a sustained benefit to auditory working memory performance and increased frontoparietal theta oscillatory power during manipulation, compared to baseline measures. These changes were not observed in participants who received sham rhTMS. This finding is important in demonstrating both behavioural and neurophysiological after-effects of non-invasive brain stimulation in a small sample of healthy individuals.

### 4.5 Limitations and future directions

Our preliminary data analyses have yielded encouraging results in a sample which, while underpowered, establishes a proof-of-concept that contributes new evidence to the cognitive neuromodulation literature. The main limitation of the current work is due to the premature suspension of data collection in compliance with public health measures at the onset of the SARS-CoV-2 pandemic. With a sample size of 7 participants per experimental group, we were not able to detect a significant between-group difference in working memory abilities between stimulation and sham control participants in the post-training session, despite finding a significant within-group improvement in d’ between pre- and post-training sessions only for the rhTMS group. Moreover this small sample size did not allow us to investigate the relationships between individual changes in behavioural performance and changes in oscillatory activity. Based on our observed effect sizes however, we can estimate the sample size necessary to draw conclusions on the behavioural after-effects of rhTMS with cognitive training, compared with cognitive training alone, which we hope will be useful for future studies.

We can estimate the effect size of our rhTMS intervention by calculating Cohen’s d for the group difference on training day 5, where we observed peak performance for the rhTMS group. Given the stimulation group mean d’ of 2.4 and the control group mean d’ of 1.4, with standard deviations of 0.9 and 1.1 respectively, Cohen’s d is 0.995, which can be interpreted as a large effect size. With this effect size estimate, the sample size would need to be 32 (17 participants per group) in order to detect an effect of stimulation, if present, using a two-tailed t-test with α = 0.05 and power = 0.8 (Faul et al., 2009).

We have demonstrated that our combined training and rhTMS protocol is feasible and tolerable for young adults without any neurologic or psychiatric conditions. Larger studies should also be appropriately powered to investigate sex-based differences in behavioural and electrophysiological results. Sex differences in working memory, particularly visuospatial rotation, have been extensively researched, and there is a large body of literature to support the association of male sex with higher proficiency in visuospatial rotation (Gurvich et al., 2020). It is plausible that biological sex is associated with neurodevelopmental differences in the robustness of connectivity within networks relevant to our cognitive training task (see Moore and Johnson (2020)) and therefore that sex may mediate response to neuromodulation, as in Weller et al. (2023). As a potential clinical tool for cognitive rehabilitation, it is imperative to ensure that our brain stimulation and training protocol is optimized to be effective for male and female patients.

In a patient population with deficits in working memory, we might expect to see different, perhaps more pronounced effects from theta rhTMS interventions, reflecting impaired functional or structural connectivity and network dysregulation (Grover et al., 2023). By the same token, we would expect improvement to be confined only to patient populations in whom the relevant white-matter pathways in the auditory dorsal stream are at least partially preserved. Identifying electrophysiological biomarkers for treatment response will also be important for identifying individual patients more likely to benefit from stimulation and training interventions (see Andrade et al. (2023)), thereby informing selection of participants for future trials and maybe one day providing personalized stimulation parameters optimized to individual network dysfunction.

## 6 Author Contributions

Conceptualization: P.A., R.J.Z. Methodology: H.T.W., P.A., R.J.Z. Investigation: H.T.W., L.K. Analysis: H.T.W., J.F.L, M.L., C.B., P.A. Resources: P.A., R.J.Z. Writing — original draft: H.T.W. Writing — review and editing: H.T.W., L.K., J.F.L., M.L., C.B., R.J.Z., P.A. Visualization: H.T.W., P.A. Supervision: R.J.Z., P.A. Project administration: P.A., R.J.Z.

## Acknowledgments

None.

