## Supplementary Material for "Information-based rhythmic transcranial magnetic stimulation accelerates learning during auditory working memory training"

### 1 Behavioural results show no evidence for cognitive transfer

#### 1.1 Auditory working memory noise task

A two-tailed Mann-Whitney U-test indicated that rhTMS and sham control groups did not significantly differ in terms of the discriminability index (d’) at pre-training (U = 18, p = 0.46) or post-training (U = 11, p = 0.10). Additionally, a one-tailed Wilcoxon signed rank test of d’ at pre-training vs. post-training, showed that neither the rhTMS group or the sham group exhibited a significant increase in performance at post-training as compared to pre-training (rhTMS W = 8, p = 0.19; sham W = 17, p = 0.71). We plotted the average d’ of both groups in both experimental sessions (Supplemental Figure 1A).

#### 1.2 Visual mental rotation task

A two-tailed Mann-Whitney U-test indicated that rhTMS and sham control groups did not significantly differ in terms of the discriminability index (d’), calculated for all angles of rotation combined, at pre-training (U = 15, p = 0.27) or post-training (U = 13, p = 0.16). However, a one-tailed Wilcoxon signed rank test of d’ at pre-training vs. post-training, showed that performance improvement at post-training as compared to pre-training for both groups (rhTMS W = 0, p = 0.02; sham W = 4, p = 0.50). We plotted the average d’ of both groups in both experimental sessions (Supplemental Figure 1B).

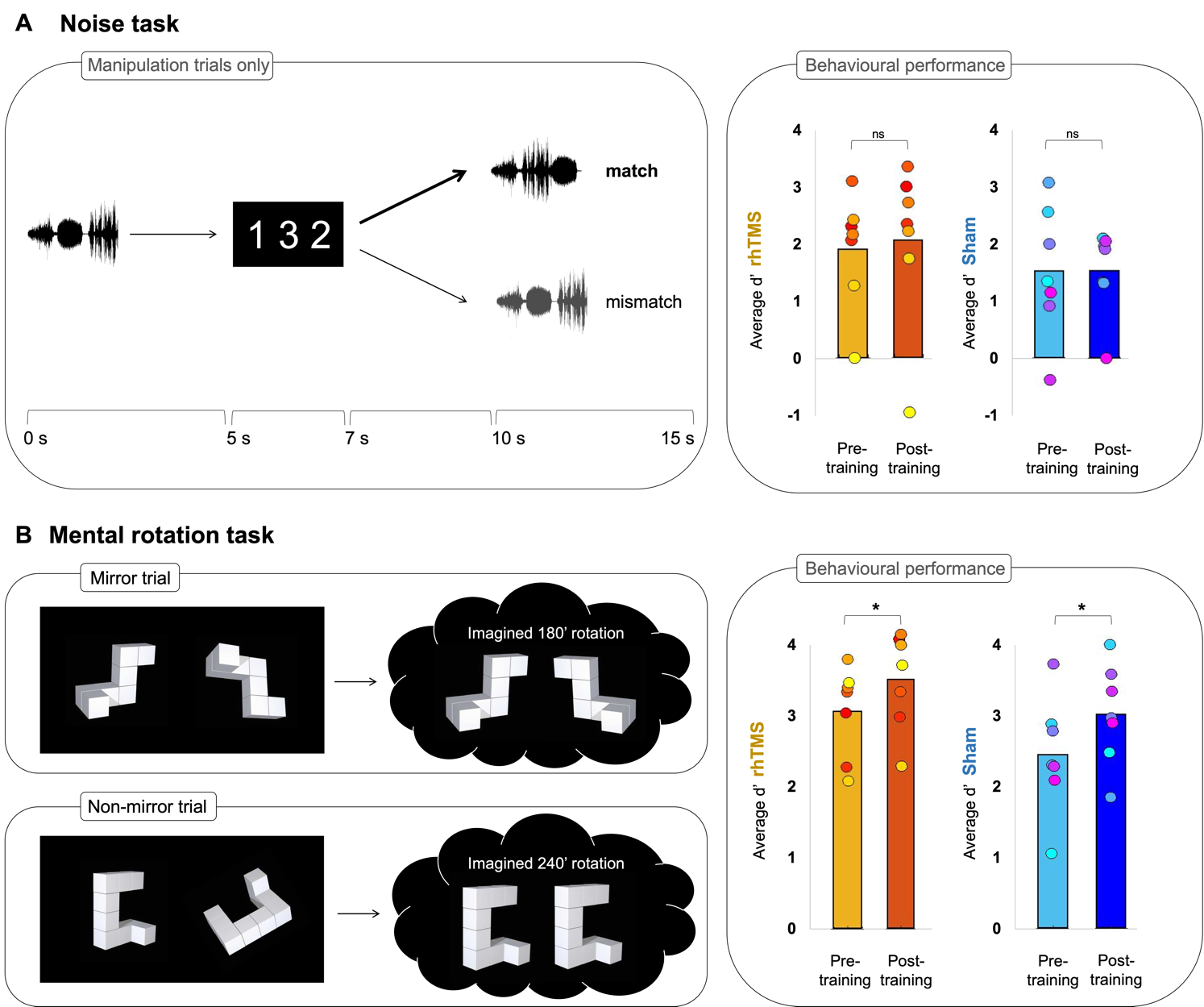

**Supplementary Figure 1: (A)** **Left panel:** Auditory working memory noise task. Timeline of a trial in the noise task as administered in pre-training and post-training sessions. Participants listened to a sequence of 3 atonal auditory stimuli. A visual string displaying the expected order of tones in the second sequence was presented 5s after the onset of the first sequence. After 5 additional seconds, participants heard another 3-tone sequence, composed of the same stimuli, and had to determine whether the order of its tones matches the order of the visual string. Correct responses are bolded for an example of a manipulation trial. There were no simple trials in this task. **Right panel:** Bar plots of the average accuracy (d’) on the noise task before and after cognitive training by experimental group; rhTMS group (n = 7, pre-training = light orange, post-training = dark orange) and sham group (n = 7, pre-training = light blue, post-training = dark blue). **(B)** **Left panel:** Visual mental rotation task. Example of a trial showing two 3D forms that are either identical (non-mirror trial) or horizontally mirrored (mirror trial). Relative to the shape on the left, the right-hand shape was rotated around its own axis by either 0, 60, 120, 180, 240, or 300 degrees. Participants had to determine whether the two forms were identical or mirror images of one another after mental rotation. **Right panel:** Bar plots of the average accuracy (d’) on the mental rotation task before and after cognitive training by experimental group; rhTMS group (n = 7, pre-training = light orange, post-training = dark orange) and sham group (n = 7, pre-training = light blue, post-training = dark blue).
